# Sudden sensory events trigger modality-independent responses across layers in the mouse neocortex

**DOI:** 10.1101/2024.11.07.622472

**Authors:** Diego Benusiglio, Hiroki Asari

## Abstract

Sudden sensory events trigger widespread electrocortical responses and noticeable changes in an animal’s behavior. It remains unclear, though, how this “surprise” signal is generated via afferent sensory input and represented by neuronal ensembles. Using in vivo electrophysiology, here we recorded the activity of different cortical areas across layers in awake head-fixed mice, while presenting sensory stimuli of different modalities and saliencies. When brief, salient stimuli were delivered at long intervals, we found prominent responses across cortical areas regardless of the stimulus modalities, resembling those observed in human electroencephalography. These responses were larger and emerged earlier in the infragranular layer than in the granular layer of the primary sensory cortex. At short inter-stimulus intervals, in contrast, only modality-specific responses were observed in the corresponding primary sensory cortex. These results indicate that sensory information travels through multiple ascending pathways, in particular, supra-modal surprise signals via non-classical sensory pathways to the cortex.

## Introduction

The percept of surprise is conserved across the animal kingdom as it serves as a major driving force for learning and adaptively adjusting behavior (Friston, 2009; Schultz, 2016; Schwartenbeck et al., 2013). Moreover, a surprising stimulus is often linked to a potential environmental threat. It is thus critical for animals to show immediate defensive responses for survival (Yilmaz and Meister, 2013), such as an increased arousal and alertness at the cognitive level, or startle reflexes at the behavioral level (Koch, 1999; Jang et al., 2015).

At the neurophysiological level, an isolated, salient and unexpected sensory stimulus evokes one of the most robust, reliable and widespread event-related potential (ERP) in human electroencephalography (EEG; Bancaud et al., 1953; Clynes, 1969; De Schoenmacker et al., 2021). This ERP is commonly referred to as vertex potential (VP) because the signal strength is maximal around the scalp vertex (Mouraux and Iannetti, 2009; Liberati et al., 2016). Growing evidence suggests that VP likely represents neural correlates of a surprise percept (Bancaud et al., 1953; Somervail et al., 2022). First, it has been recorded not only in humans but widely across species, including rats and non-human primates (Somervail et al., 2021; November et al., 2024). Second, the magnitude of VP diminishes if the stimulus becomes no longer surprising; e.g., when the same stimulus is presented repeatedly in regular intervals, hence more predictable (Ronga et al., 2013). Third, VP can be triggered regardless of the stimulus modality (Mouraux and Iannetti, 2009; Liberati et al., 2016). Thus, VP does not simply reflect modality-specific sensory processing mediated by the classical sensory pathways, but rather represents a supra-modal signature of the stimulus, such as surprise, presumably via non-classical sensory pathways (Guillery, 2005; Bastos et al., 2012). It remains unclear, though, how such sensory signals reach the cortex and form a neural representation of a surprise percept.

To address this question, here we took an in vivo electrophysiology approach to probe the mouse cortical responses to sudden sensory stimuli. In particular, we designed our stimulus protocol in accordance with previous human studies (Braff et al., 2001; Grill-Spector et al., 2006; Mancini et al., 2018), and identified homologous ERP characteristics in the mouse EEG to the human VP. Furthermore, intracortical recordings in awake head-fixed mice revealed that a) sudden sensory stimuli drove the infra-and supra-granular layers of the primary sensory cortex more strongly than the granular layer, regardless of the stimulus modalities; and b) the infragranular layer was driven before the granular layer. These results support that information about a sensory surprise is transmitted to the mouse neocortex via non-classical sensory pathways.

## Results

### Abrupt sensory stimulation triggers vertex potential homologues in the mouse cortex

To investigate the mouse cortical activity in response to sudden sensory events, we first monitored the electrical activity at the macroscopic level in awake head-fixed mice (**Fig. 1A**). In particular, we recorded electroencephalography (EEG) signals using a low-impedance micro-screw electrode attached to the skull (AP -1.0 mm, ML -1.5 mm, DV 0.0 mm from Bregma), and presented abrupt stimuli of three different modalities (auditory, 8 kHz pure tone; somatosensory, electrical tail-shock; visual, full-field flash) at various inter-stimulus intervals (ISIs; **Fig. 1B**). The recorded EEG signal was then aligned to the stimulus onset, and averaged across trials to estimate the event-related potentials (ERPs) for each sensory modality separately.

**Figure 1:**
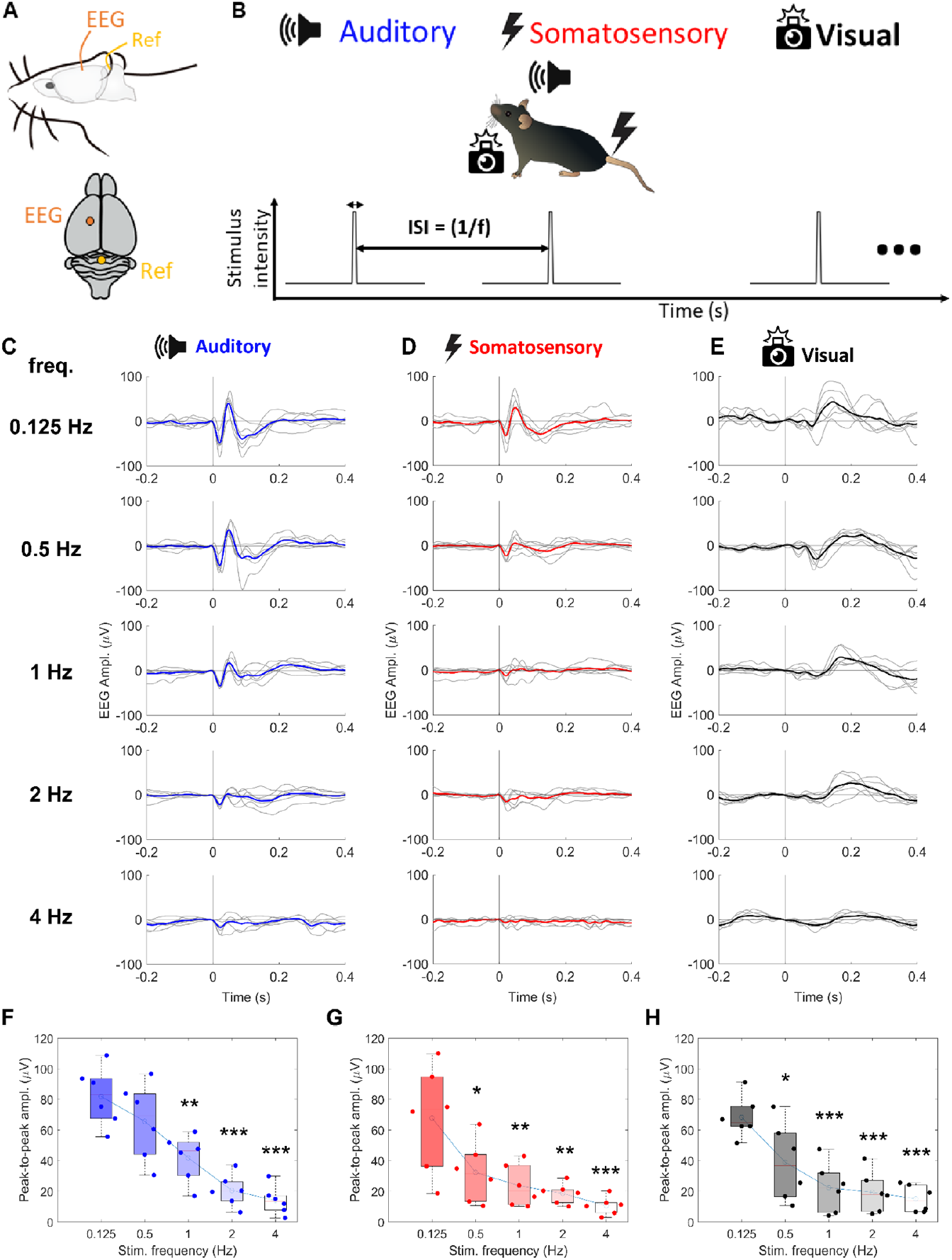
The mouse electroencephalogram displays homologues of the human vertex potential in response to sudden sensory events. *A*: Schematic diagram of chronic EEG recordings in awake head-fixed mice for the ERP analysis evoked by sudden sensory events. *B*: Schematic diagram of sensory stimulation paradigm, commonly used across three different modalities (blue, auditory; red, somatosensory; black, visual stimulation). *C*: Auditory ERPs at five different temporal frequencies (from top to bottom: 0.125, 0.5, 1, 2, and 4 Hz): thin gray lines, trial-average (40 trials for each condition) for each mouse; thick colored line, grand-average across mice (n = 6 animals). *D,E*: Corresponding figure panels for somatosensory and visual ERPs, respectively. *F-G*: Boxplot of peak-to-peak amplitudes of ERPs at different temporal frequency stimulations (F, auditory; G, somatosensory; H, visual stimuli). One-way analysis-of-variance (ANOVA) with Tukey-Kramer test against 0.125 Hz stimulation data: *, p < 0.05; **, p < 0.01; ***, p < 0.001. See **Suppl. Table 1** for post-hoc test p-values of all possible frequency pairs.

**Supplementary Table 1:** Post-hoc test p-values of all possible frequency pairs, related to Fig. 1F-H.

| Stim. Freq. (Hz) | vs. stim. freq. (Hz) | corrected p-values |  |  |
| --- | --- | --- | --- | --- |
|  |  | Auditory | Somatosensory | Visual |
| 0.125 | 0.5 | 0.477 | 0.029 (*) | 0.036 (*) |
| 0.125 | 1 | 0.003 (**) | 0.0044 (**) | 0.00051 (***) |
| 0.125 | 2 | 1.43e-05 (***) | 0.0015 (**) | 0.00023 (***) |
| 0.125 | 4 | 2.84e-06 (***) | 0.00023 (***) | 7.34e-05 (***) |
| 0.5 | 1 | 0.141 | 0.932 | 0.428 |
| 0.5 | 2 | 0.00094 (***) | 0.732 | 0.268 |
| 0.5 | 4 | 0.00017 (***) | 0.314 | 0.121 |
| 1 | 2 | 0.226 | 0.991 | 0.998 |
| 1 | 4 | 0.0632 | 0.766 | 0.938 |
| 2 | 4 | 0.962 | 0.948 | 0.991 |

Following the experimental design in previous human studies (Braff et al., 2001; Grill-Spector et al., 2006; Mancini et al., 2018), we repetitively presented the sensory stimuli of each modality at five different temporal frequencies: 0.125, 0.5, 1, 2, and 4 Hz. The stimulus frequencies and modalities were randomized across block presentations. We found larger ERPs for lower temporal frequencies (i.e., longer ISIs; **Fig. 1C-H**), while the first peak latencies of those ERPs were largely unaffected across all stimulus modalities (**Suppl. Fig. 1**). In particular, stimulation at the lowest temporal frequency (0.125 Hz, or 8 s ISI) robustly evoked a pronounced ERP with similar dynamics across modalities and trials (auditory, 81.9 ± 7.9 µV; somatosensory, 67.8 ± 14.1; visual, 68.3 ± 5.6; mean peak-to-peak ERP amplitude ± standard error, n=6 animals). Although the mice sometimes displayed a motor reaction to the onset of such sensory events, the movement was not positively correlated with the ERPs at the single-trial level (**Suppl. Fig. 1**). These results suggest that the observed supra-modal ERPs likely represent a mouse homologue of the human vertex potential (VP), a putative neural correlate of sensory surprise (Mouraux and Iannetti, 2009).

**Supplementary Figure 1:**
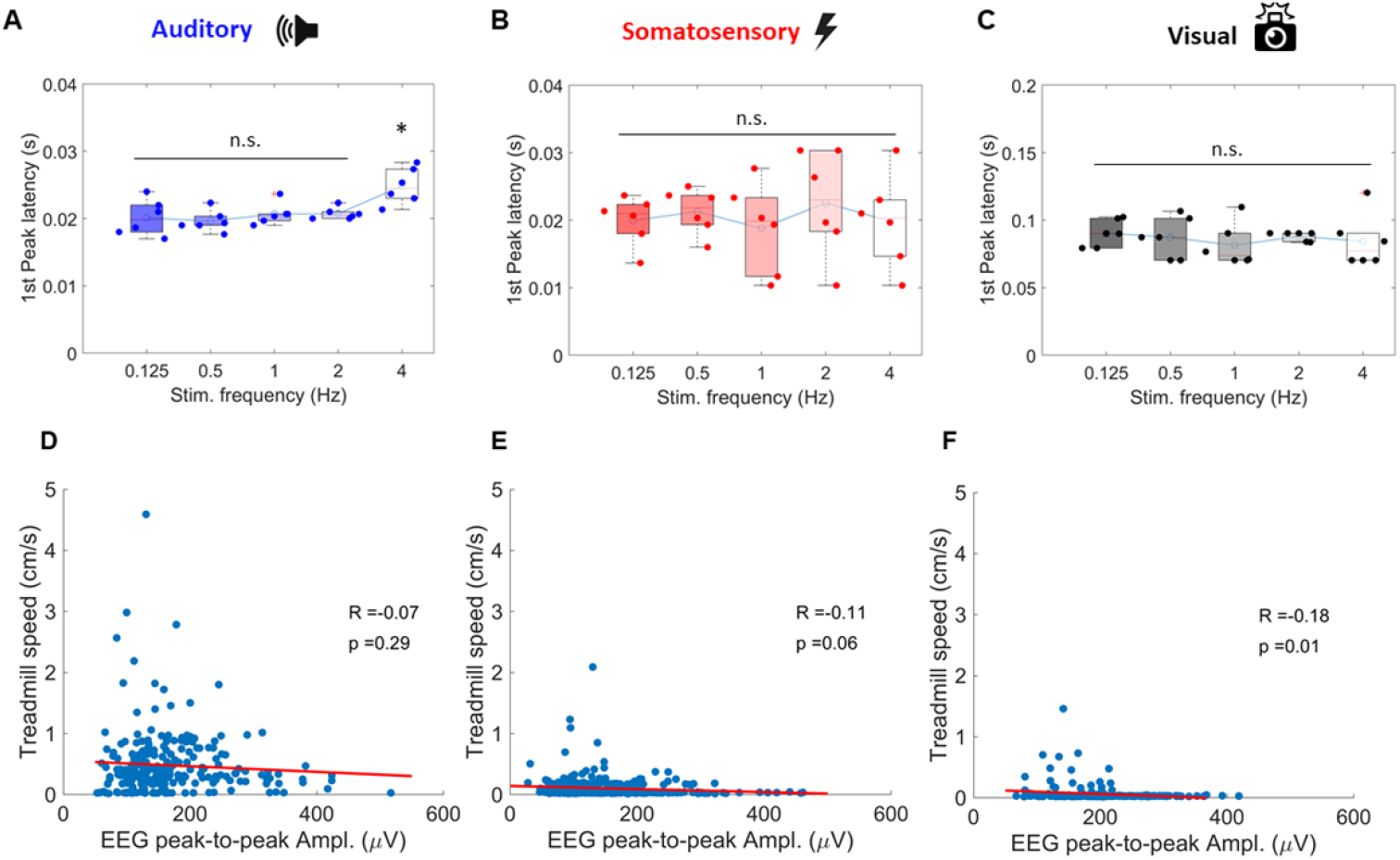
The dynamics of the mouse vertex potential homologues are not affected by the stimulation temporal frequencies or the animal’s movement. *A*: Boxplot of the first peak latencies of auditory event-related potentials (ERPs) at five different stimulation frequencies (0.125, 0.5, 1, 2, and 4 Hz; n = 6 animals). One-way analysis-of-variance with Tukey-Kramer test against the 0.125 Hz stimulation condition: *, p < 0.05; **, p < 0.01; *** p < 0.001. We found p > 0.05 in all pairwise comparisons except for 4 Hz versus 0.125 Hz (p = 0.03). *B,C*: Corresponding figure panels for somatosensory and visual ERPs, respectively. *D-F*: Pearson correlation coefficient between the magnitude of the sensory-evoked ERPs (D, auditory; E, somatosensory; F, visual; 40 trials for each condition for each animal) and the simultaneously recorded movements of the animal (n = 6 animals). We found no – or rather negative, if any – correlation, supporting that the sudden sensory stimuli used in this study were at the sub-startle level.

To further examine a similarity to the human VP, we next investigated how the mouse ERPs exhibit short-term adaptation to repetitive stimulus presentations. In particular, we presented triplets of stimuli (ISI, 1 s) at an interval of 6-10 s between the triplets (drawn from a random uniform distribution), and found a significant reduction of the ERP magnitude already at the second stimulus of the triplet, compared to the first one (**Suppl. Fig. 2**). This rapid habituation of the mouse ERP is consistent with the one observed in the human VP (Mancini et al., 2018), further supporting the homology of the cortical sensory surprise responses across species (Somervail et al., 2021; November et al., 2024).

**Supplementary Figure 2:**
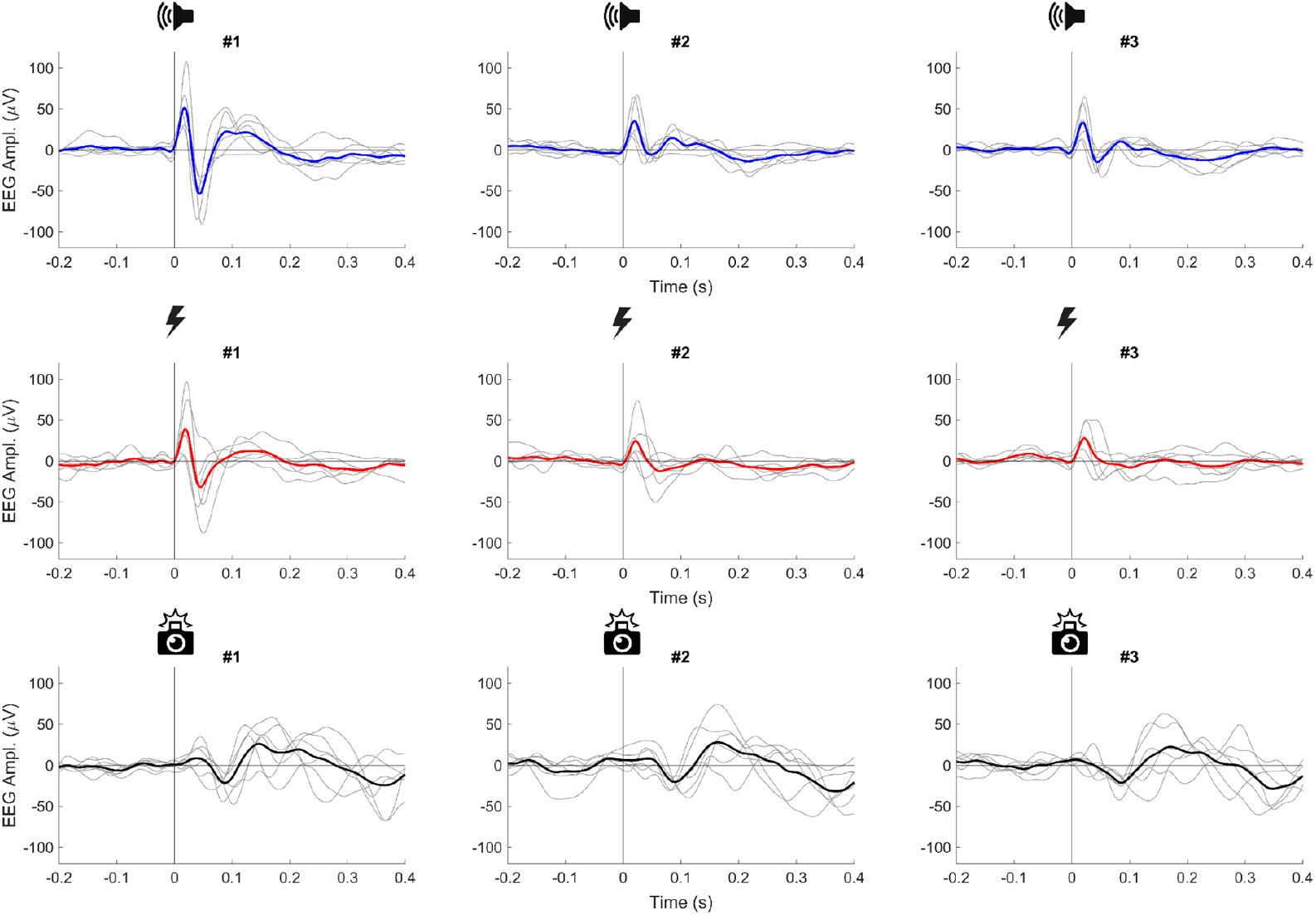
The mouse vertex potential homologues show rapid adaptation to repeated stimulations. Average ERPs evoked by triplets of stimuli at 1 s inter-stimulus intervals for auditory (top, blue), somatosensory (middle, red), and visual (bottom, black) modalities (inter-triplet intervals, 6-10 s drawn from a random uniform distribution). Thin gray lines, trial-average (40 trials for each condition) for each mouse; thick colored line, grand-average across mice (n = 6 animals). Average peak-to-peak amplitude of the ERP decreased substantially from the 1st stimulation (left) to the 2nd (middle) and 3rd (right) ones: auditory, 114.7, 50.3, 53.7 µV; somatosensory, 80.9, 45.4, 40.8 µV; visual, 64.1, 54.5, 50.8 µV.

### Abrupt sensory stimulation drives the mouse cortex via non-classical sensory pathways

How does the mouse VP homologue arise? Sensory signals travel from the peripheral sensory organs to the mammalian neocortex via multiple ascending pathways (Mountcastle, 1980; Heimer et al., 1983). For example, the so-called classical sensory pathway, such as the dorsal column-medial lemniscus of the somatosensory system, transmits modality-specific environmental information to the corresponding primary sensory cortex, and is considered to faithfully encode physical properties of sensory stimuli, such as their intensity and frequency (Chapin and Lin, 1984; Seelke et al., 2012). In contrast, non-classical pathways are less specific to a given sensory modality, but receive and integrate inputs from the other sensory systems. They are thus considered to represent multi-modal information, including sensory surprise (Jones et al., 2001; Henschke et al., 2015).

To better understand the encoding of sensory surprise in the mouse brain, we next examined the primary sensory cortical responses to sudden stimuli of the congruent and non-congruent modalities. Specifically, using multichannel silicon probes, we recorded the local field potential (LFP) and multi-unit spiking activity across layers of the primary somatosensory cortex (S1) or the primary visual cortex (V1), while presenting either somatosensory, visual, or auditory stimuli at five temporal frequencies as we did with the EEG recordings (**Fig. 1B**).

In response to non-congruent modality stimuli at low temporal frequencies, we found prominent responses across layers in the primary sensory cortex (**Fig. 2A**). In particular, the total transmembrane current flow (TTCF) on the current source density (CSD) demonstrated larger responses in the infragranular (layer V/VI) and the supragranular (layer II/III) layers than in the granular layer (layer IV; **Fig. 2B**). The infragranular layer responded first (**Fig. 2C**), and the latency of the CSD responses was compatible with that of the EEG responses (∼20 ms for auditory and somatosensory ERPs and ∼75 ms for visual ERP; **Fig. 1**). In contrast, a high frequency stimulation triggered no substantial response in the primary sensory cortex of non-congruent modalities, much as in the EEG recordings. These results suggest that information about abrupt sensory events is processed via non-classical sensory pathways.

**Figure 2:**
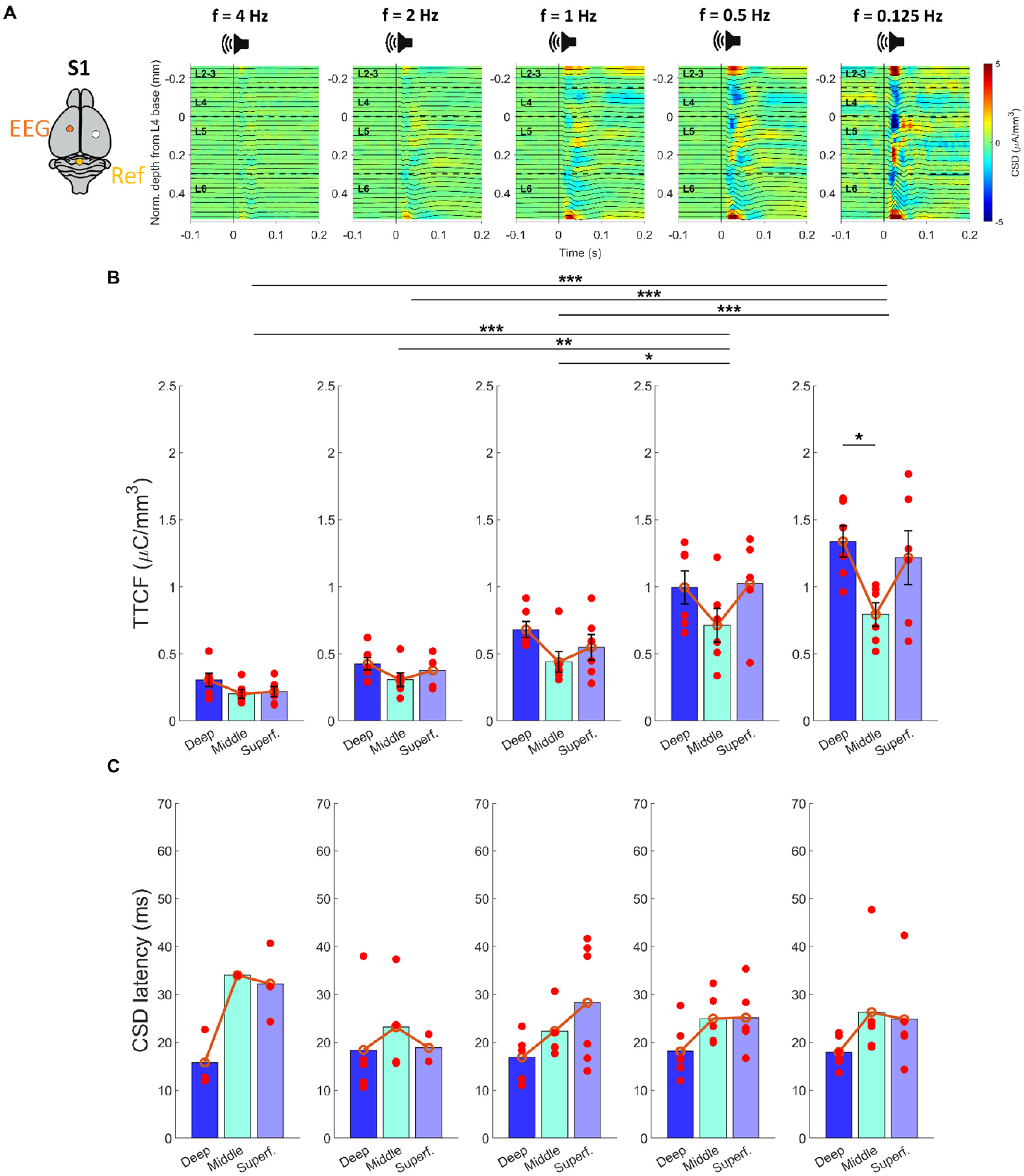
Sudden sensory stimuli drive the primary sensory cortex of non-congruent modalities across layers via the infragranular layer. *A*: Representative CSD (heatmap) and LFP (overlaid lines) in S1 in response to brief, loud pure tones (8 kHz, 50 ms, 80 dB SPL) at five different temporal frequencies (from left to right: 4, 2, 1, 0.5, and 0.125 Hz). *B*: Total transmembrane current flow (TTCF) integrated over 0-70 ms post-stimulus window at different recording depths (deep infragranular, middle granular, and superficial supragranular cortical layers) for each stimulation frequency (trial average, N=6 animals). *C*: Latency of the CSD response onset at different cortical layers for each stimulation frequency. See **Suppl. Table 2** for post-hoc test p-values of all possible frequency pairs.

**Supplementary Table 2:** Post-hoc test p-values of all possible frequency pairs, related to Fig. 2B,C.

| Stim. Freq. (Hz) | vs. stim. freq. (Hz) | corrected p-values |  |
| --- | --- | --- | --- |
|  |  | TTCF | CSD latency |
| 0.125 | 0.5 | 0.59 | 0.99 |
| 0.125 | 1 | 5.4 e-04 (***) | 0.97 |
| 0.125 | 2 | 3.1 e-05 (***) | 0.99 |
| 0.125 | 4 | 8.7 e-07 (***) | 0.99 |
| 0.5 | 1 | 0.03 (*) | 0.95 |
| 0.5 | 2 | 0.002 (**) | 0.98 |
| 0.5 | 4 | 5.4 e -05 (***) | 0.99 |
| 1 | 2 | 0.63 | 0.99 |
| 1 | 4 | 0.08 | 0.95 |
| 2 | 4 | 0.74 | 0.97 |

We observed distinct activity patterns in the mouse primary sensory cortex when stimuli of the congruent sensory modality were presented (**Fig. 3**). First, the modality-specific cortical responses remained salient even at higher temporal frequency stimulations (e.g., 4 Hz; **Fig. 3A,B**). The granular layer generally showed the strongest responses at all temporal frequencies we examined, except for the lowest frequency stimulation. At 0.125 Hz, the infragranular layer showed stronger responses than the granular layer, much as those responses evoked by the non-congruent modality stimuli (**Fig. 2**). Second, the latencies of the cortical responses were generally shorter for the congruent modality stimuli than for the non-congruent ones (**Fig. 3C**). These results indicate that, as expected, modality-specific sensory signals travel via the classical sensory pathway to reach the corresponding primary sensory cortex.

**Figure 3:**
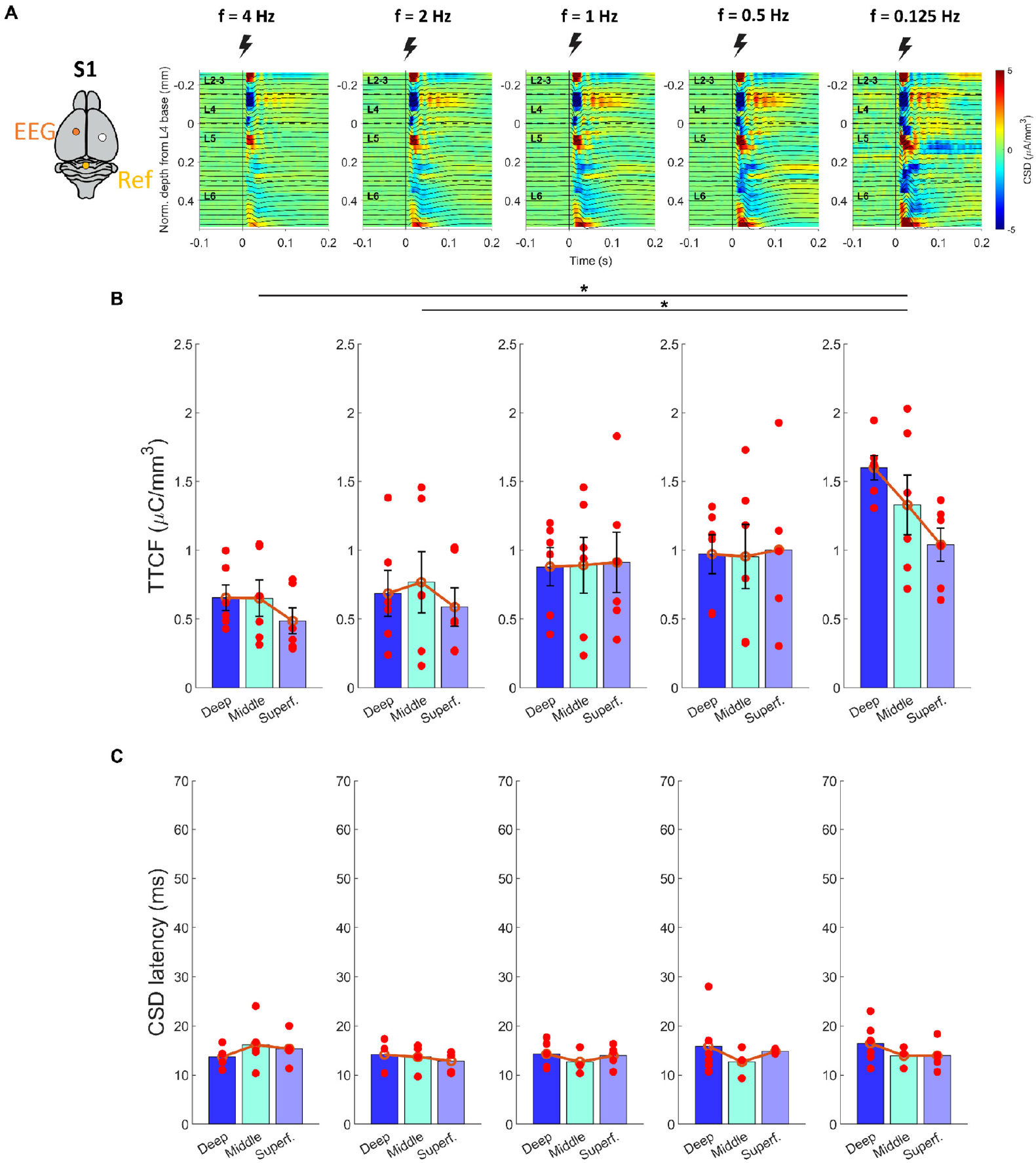
The granular layer of the mouse primary sensory cortex remains responsive to repetitive presentations of the congruent modality stimuli. *A*: Representative CSD (heatmap) and LFP (overlaid lines) in S1 in response to brief, electrical tail-stimulation at five different temporal frequencies (from left to right: 4, 2, 1, 0.5, and 0.125 Hz). *B, C*: TTCF (B: 0-70 ms post-stimulation window, averaged across trials; N=6 animals) and latency of the CSD response onset (C) at different recording depths for each stimulation frequency. Figure format follows that in **Fig.2**. See **Suppl. Table 3** for post-hoc test p-values of all possible frequency pairs.

**Supplementary Table 3:** Post-hoc test p-values of all possible frequency pairs, related to Fig. 3B,C.

| Stim. Freq. (Hz) | vs. stim. freq. (Hz) | corrected p-values |  |
| --- | --- | --- | --- |
|  |  | TTCF | CSD latency |
| 0.125 | 0.5 | 0.30 | 0.99 |
| 0.125 | 1 | 0.25 | 0.84 |
| 0.125 | 2 | 0.03 (*) | 0.79 |
| 0.125 | 4 | 0.01 (*) | 0.99 |
| 0.5 | 1 | 0.99 | 0.97 |
| 0.5 | 2 | 0.83 | 0.96 |
| 0.5 | 4 | 0.61 | 0.99 |
| 1 | 2 | 0.83 | 0.97 |
| 1 | 4 | 0.60 | 0.96 |
| 2 | 4 | 0.99 | 0.95 |

## Discussion

Sensory surprise is an innate perception that arises when an animal encounters an isolated, salient and unexpected sensory stimulus of any modality. To study how sensory surprise is encoded in the mouse brain, here we recorded the cortical responses to brief sensory stimuli of three different modalities, each at five different inter-stimulus intervals, in awake head-fixed mice (**Fig. 1**). As expected, when sensory stimuli were repetitively presented at a high temporal frequency, the granular layer of the primary sensory cortex of the same modality was driven first and most prominently, suggesting that the signal reached there via the classical ascending sensory pathway (**Fig. 3**). In contrast, at a low temporal frequency, we found that sudden sensory events triggered robust responses widely across cortical areas and layers, independent of the sensory modalities (**Figs. 1-3**), similar to the so-called vertex potential (VP) observed in human electroencephalography (EEG; Mouraux and Iannetti, 2009). In particular, the infra- and supra-granular layers were evoked before the granular layer (**Fig. 2**), suggesting that the signal reached via non-classical sensory pathways.

What does this mouse homologue of the human VP represent? We consider it as a neurophysiological signature of a sensory surprise percept for the following reasons. First, we observed larger cortical responses when the stimulus was presented with longer inter-stimulus intervals, causing a stronger violation of quiescence, regardless of the stimulus modalities (**Fig. 1**). This is consistent with the definition that a sensory surprise is triggered by an unexpected event that violates prior expectations or predictions (Friston, 2009). Second, no valence was associated with the stimuli presented in this study. Sensory surprise is linked to the concept of prediction errors, and was extensively studied in the context of reinforcement learning (Schultz, 2016). However, here we presented sensory stimuli to passively receiving animals without imposing any learning tasks. The stimuli did not carry any positive or negative valence, hence presumably inducing a neutral percept of sensory surprise. Third, we barely observed reflexive motor responses of the animals during the experiments (**Suppl. Fig. 1D-F**) as we used sub-startle sensory stimuli: e.g., around 80 dB sound pressure level (SPL) for auditory stimuli, while acoustic startle responses in mice require >95 dB SPL stimulation (Pantoni et al., 2020). Furthermore, we found no positive correlation between the evoked neuronal signals and the movement of the animals. Thus, while sensory surprise often accompanies a startle response as an immediate defensive reaction, the neuronal signals we observed have little to do with such behavior. Lastly, we used repetitive stimulations of an identical brief stimulus at a regular interval in each sensory modality block. Thus, the average responses we analyzed here were irrelevant to the familiarity of the stimulus itself, hence different from a novelty response or mismatch negativity (Chen et al., 2015; Ross and Hamm, 2020). Taken together, we think that the observed signals largely represent neuronal correlates of sensory surprise.

In conclusion, here we demonstrated the mouse homologue of the human vertex potential in response to sudden sensory stimuli, and such sensory information was transmitted to the mouse neocortex via non-classical sensory pathways. A surprising stimulus is known to induce substantial changes in the brain states (Antony et al., 2020; English et al., 2023). It is a future challenge to study how this cortex-wide surge of the activity interacts with sensory information conveyed from the classic sensory pathways and eventually affects the behavior of an animal.

## Materials and Methods

No statistical method was used to predetermine the sample size. Experimental procedures involving animals were performed under the license 233/2017-PR and 220/2024-PR from the Italian Ministry of Health, following protocols approved by the Institutional Animal Care and Use Committee at European Molecular Biology Laboratory. The data analyses were done in Python 3 and Matlab R2021b-2023a (Mathworks). The statistical significance level was set to be 0.05. All summary statistics were described as the mean and the standard error of the mean unless otherwise stated.

### Animals

Twelve C57BL/6J male mice were used at 8-9 weeks of age at the time of the surgery. The mice were housed under standard laboratory conditions (12-hour-light / 12-hour-dark cycle, 20-24 °C, 55 ± 10 % humidity, water and food *ad libitum*). After the implantation of a head-plate for electrophysiological recordings, the animals were kept single-housed.

### Surgery

We performed head-plate implantation as described previously (Boissonnet et al., 2023). In brief, the animal was weighed and injected intraperitoneally with a mixed solution of antibiotic (Baytril, Bayer, 5 mg/kg) and analgesic (Rimadyl, Zoetis, 5 mg/kg). The animal was then anesthetized (induction, 4% isoflurane in oxygen; maintenance, 1.5-2%) and placed inside a stereotaxic apparatus (Stoelting 51625). Throughout the surgical procedure, body temperature was maintained at 37°C using a heating pad (Supertech Physiological), and eye ointment (VitA-POS) was used to prevent the eyes from drying. After positioning the head using the ear-bars, the scalp skin was disinfected (Betadine) and removed with scissors. Soft tissue was removed from the skull surface with a round scalpel, and ethanol and acetone were applied to the skull to disinfect and remove any residual compounds. Tissue adhesive (Vetbond, 3M) was used to fix and seal the skin edges. A custom-made titanium head-plate (0.8 mm thick) was then cemented to the skull using acrylic cement powder (Paladur, Kulzer) pre-mixed with cyanoacrylate adhesive (Loctite 401, Henkel). A well was made with dental cement around the marks later to hold saline solution over the brain. Two gold plated beryllium copper micro-screws (0.47 mm, Preci-dip) were fixed onto the skull at the following stereotaxic coordinates from Bregma: 1) AP: -1.00 mm, ML: -1.50 mm; 2) AP: -5.80 mm, ML: 0.00 mm. The micro-screws were later used during the recording sessions as electroencephalography (EEG) electrodes.

After the surgical procedures, the animals were recovered from anesthesia in a warmed-up chamber and returned to their home cage. For postoperative care, the animals were given intraperitoneally antibiotic and analgesic daily for 3-5 days and health status was checked daily for 7 days. We waited another 10 days until the cranial window completely recovered before starting acute electrophysiological recording sessions.

### In vivo electrophysiology

The day before the electrophysiological recording session, the animal was anesthetized and placed inside a stereotaxic apparatus with a heating pad. The scalp was registered in the stereotaxic controller, and entry points for targeting the primary somatosensory cortex (S1; AP: -1.50 mm, ML: 2.00 mm) or primary visual cortex (V1; AP: -3.80 mm, ML: 2.00 mm) were marked with a high-speed surgical drill (OmniDrill 35, WPI). A craniotomy (diameter, 1 mm) was made and soaked with sterile cortex buffer (NaCl 125 mM, KCl 5 mM, Glucose 10 mM, HEPES 10 mM, CaCl_2_ 2 mM, MgSO_4_ 2 mM, pH 7.4). A protective silicon elastomer plug (Kwik-Sil, World Precision Instruments) was placed on top of the skull to protect the craniotomy.

On the day of the recording session, the awake animal was placed on a circular treadmill with its head fixed, and the silicon plug was removed to expose the craniotomy that was kept soaked with a sterile cortex buffer. An acute silicone probe with 32 recording contacts (Assy-37, Cambridge Neurotech) coated with a fluorescent dye (DiI stain, Invitrogen, D282) was slowly inserted into the brain (100 μm/min) with a micromanipulator (Neurostar) attached to the stereotaxic apparatus. The probe was moved until cortical activity was visible as rhythmic fluctuations of local field potential (LFP) across all electrodes. The LPF and action potentials of cortical neurons were monitored online during the procedure with a visualization software (Smartbox Interface, NeuroNexus). The electroencephalography (EEG) signal, measured as the difference of the electrical potential between the two micro-screws, was amplified by an analogue amplifier (Model 1700, KF Technology) with hardware band-pass filter 1-20 kHz. The electrophysiology data, the treadmill sensor, and the sensory stimulation trigger signals were recorded at 30 kHz per channel (SmartBox, NeuroNexus). The locomotion of the animal, its body posture and pupil size were monitored at 60 Hz using cameras (DMK33UX174, Imaging Source) under infrared illumination.

At the end of the recording session, the electrode location was verified histologically (**Suppl. Fig. 3**). After retracting the silicone probe, the animal was anesthetized (2.5% Avertin, 16 μL/g, intraperitoneal injection) and perfused with paraformaldehyde (PFA; 4% in phosphate buffer solution). The brain tissue was harvested and post-fixed overnight in 4% PFA at 4°C. Coronal sections of the brain tissue (thickness, 100 μm) were then examined under a fluorescence microscope (Leica, LMD7000 with N2.1 filter cube) to visualize the trace left by the DiI stain on the probe.

**Supplementary Figure 3:**
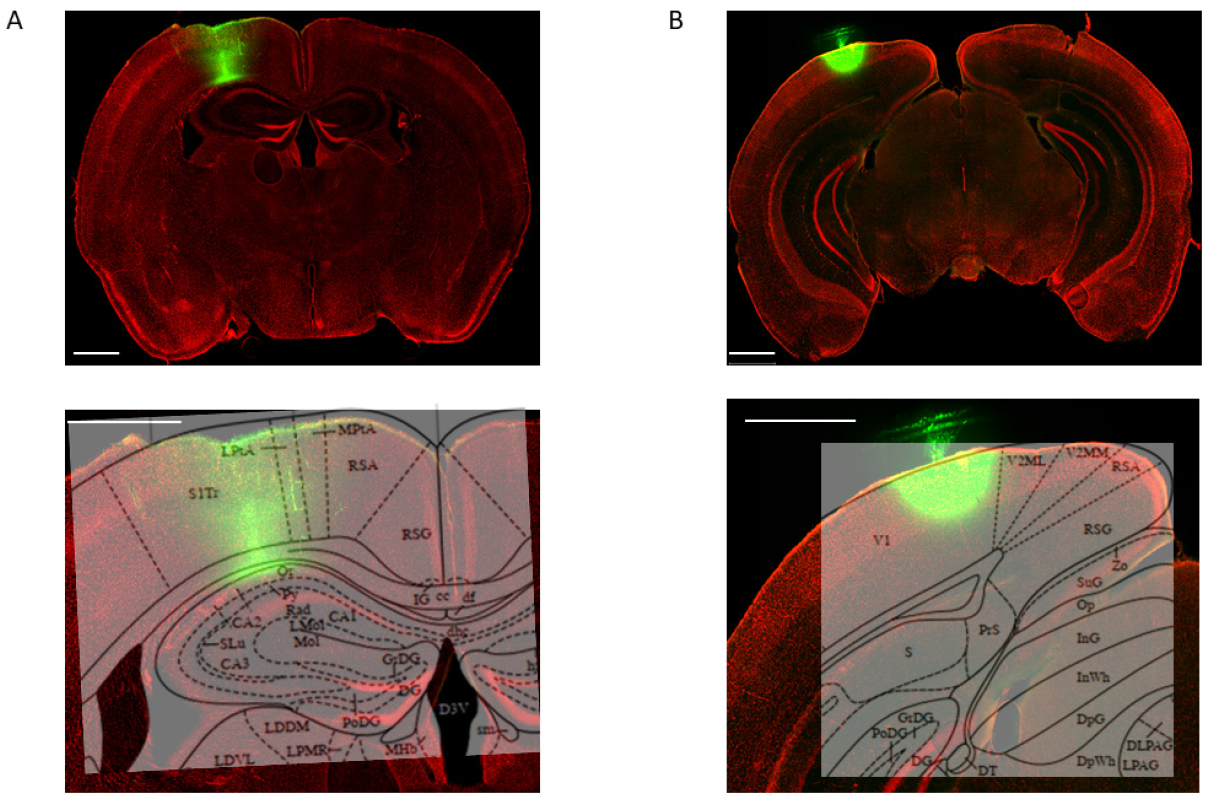
Histological examination of the intracortical electrode location. *A*: Top, representative image of the coronal brain slice (DAPI staining, red) with the trace of the DiI-coated probe (green) in the mouse S1. Bottom, magnified image with the overlaid reference mouse brain atlas (Paxinos and Franklin, 2001). Scale bar, 1 mm in all images. *B*: representative image of the probe trace in V1 (shown in the same format as panel A).

### Sensory stimulation

For auditory stimuli, we used brief, fast-rising tones (50-ms duration, 5-ms rise and fall times, frequency 8 kHz) presented at around 80 dB sound pressure level (SPL), delivered through a speaker (stereo speaker, Trust) placed in front of the mouse at around 50 cm of distance.

For somatosensory stimuli, we delivered constant-current biphasic square-wave electrical pulses (0.4-0.8 mA, 200-µs duration, Isostim A320, World Precision Instruments) to the mouse tail using bipolar copper clamp electrodes (6 mm, Mueller electric). The intensity of somatosensory stimulation was adjusted for each mouse to elicit a reproducible tail twitch, but not any reflexive body reaction.

Visual stimuli were presented as described before (Boissonnet et al., 2023). Briefly, a gamma-corrected digital light processing device (DLP, Texas Instruments, DLPDLCR3010EVM-LC; green and red light-emitting diodes replaced with ultraviolet and infrared ones, respectively) was used as a light source to project visual images onto a spherical screen (20 cm in radius), placed 20 cm from the mouse eye. Full-field flashes (50 ms duration) were presented at the maximum intensity of 31 mW/m^2^ at the mouse eye position.

### Stimulus design

Stimuli of the three sensory modalities (auditory, somatosensory, and visual) were presented in blocks of 40 trials, respectively, at five different interstimulus intervals (ISIs), resulting in the following temporal frequencies: 0.125, 0.5, 1, 2, and 4 Hz. The order of stimulus frequencies and modalities was randomized across block presentations. Block of 40 triplets of stimuli (1 s ISI) of the same modality were presented at an inter-triplet interval of 6-10 s (drawn from a uniform random distribution).

## Data analysis

### Electroencephalography and event-related potential analysis

The recorded EEG signals were band-pass filtered (0.5-30 Hz, Butterworth filter, 3^rd^ order), segmented into 2-s long epochs relative to the onset of each sensory stimulus (from -1 s to +1 s), baseline corrected in the pre-stimulus window (−0.01 s to -0.002 s), and averaged across trials to identify the event-related potentials (ERPs). As a feature of the ERP, we focused on the peak-to-peak amplitude (e.g., **Fig. 1C-H**) and the first peak latency (e.g., **Suppl. Fig. 1A-C**), and performed analysis-of-variance (ANOVA) analysis for statistical evaluations (**Suppl. Table 1**).

### Local field potentials and current source density analysis

The extracellular LFP data were band-pass filtered (0.5-100 Hz, Butterworth filter, 3^rd^ order), segmented into 2-s long epochs relative to the onset of each sensory stimulus (from -1 s to +1 s), baseline corrected in the pre-stimulus window (−0.01 s to -0.002 s), and averaged across trials to identify the ERPs.

To estimate cortical depth of the electrode, we computed the current source density (CSD) and analyzed the sensory-evoked source/sink distributions for the congruent sensory modality (somatosensory stimuli in S1 and visual stimuli in V1; e.g., **Figs. 2A and 3A**). The CSD was computed on the average of 40 stimuli. The earliest onset current sink visible across at least three neighboring channels was identified as putative granular layer (cortical layer IV), as described previously (De Franceschi, et al., 2011). A consensus on the estimation of the putative granular layer was made by comparing the CSD of the congruent sensory modality for all stimuli frequency. The location of the granular layer was also confirmed with the corresponding histological images (**Suppl. Fig. 3**).

To evaluate the overall magnitude of the stimulus-evoked currents in the cortex, we first calculated the total transmembrane current flow (TTCF) by integrating the rectified CSD signals over the 0-70 ms post-stimulus time window (**Figs. 2B and 3B**). We then averaged the TTCF across those channels in each of the three cortical depth categories: superficial (cortical layer II-III), middle (cortical layer IV), and deep (cortical layer V-VI). The latencies of the CSD responses were calculated as the earliest time (onset) at which the rectified CSD exceeded the threshold of 2 standard deviations of the CSD calculated in the 2-s long epochs. The latency of the CSD responses were then identified with the average signal across channels in each of the three cortical depth categories (**Figs. 2C and 3C**). We performed ANOVA analysis for comparing these measures across cortical depths and stimulation temporal frequencies (**Suppl. Tables 2 and 3**).

## Acronyms

CSD: current source density
EEG: electroencephalography
ERP: event-related potential
ISI: inter-stimulus interval
LFP: local field potential
TTCF: total transmembrane current flow
VP: vertex potential.

## Acknowledgements

This work was supported by research grants from EMBL (H.A.); and EMBL-IIT Postdoctoral Fellowship (ETPOD; D.B.). EMBL IT Support is acknowledged for provision of computer and data storage servers; and the LAR facility for taking care of animals. We thank all the Asari lab members as well as Gian Domenico Iannetti (Italian Institute of Technology) and his group members for many useful discussions.

